# Reference transcriptome data in silkworm *Bombyx mori*

**DOI:** 10.1101/805978

**Authors:** Kakeru Yokoi, Takuya Tsubota, Akiya Jouraku, Hideki Sezutsu, Hidemasa Bono

**Author notes:** These authors equally contributed to this work (Kakeru Yokoi, Takuya Tsubota). Corresponding author: Kakeru Yokoi. E-mail addresses: Takuya Tsubota, Akiya Jouraku, Hideki Sezutsu, Hidemasa Bono.

## Abstract

**Background:** The silkworm *Bombyx mori* is a lepidopteran model insect with biological and industrial importance. Its high-quality genome sequence has recently become available and the utilization of this information in combination with extensive transcriptomic analyses is expected to provide an elaborate gene model. It will also be possible to clarify gene expression in detail using this approach.

**Results:** We herein performed RNA-seq analysis of ten major tissues/subparts of silkworm larvae. Sequences were mapped onto the reference genome assembly and reference transcriptome data was successfully constructed. The reference data provided a nearly complete sequence for *sericin*-*1*, a major silk gene with a complex structure. We also markedly improved the gene model for other genes. Transcriptomic expression was investigated in each tissue and a number of transcripts were identified that were exclusively expressed in tissues such as the testis. Transcripts strongly expressed in the midgut formed tight genomic clusters, suggesting that they originated from tandem gene duplication. Transcriptional factor genes expressed in specific tissues or the silk gland subparts were also identified.

**Conclusions:** We successfully constructed reference transcriptome data in the silkworm and found that a number of transcripts showed unique expression profiles. These results will facilitate basic studies on the silkworm and accelerate its applications, which will contribute to further advances in lepidopteran and entomological research and the practical use of these insects.

## Background

The silkworm *Bombyx mori* is a lepidopteran insect that has been utilized in studies of physiology, genetics, molecular biology, and pathology. Functional analyses of genes related to hormone synthesis/degradation, pheromone reception, larval marking formation, and virus resistance have been performed using this silkworm [1–5], and the findings obtained have contributed to the promotion of insect science. The silkworm has the ability to produce large amounts of silk proteins, which is one of the most prominent characteristics of this species. Silk proteins are mainly composed of the fibrous protein Fibroin and aqueous protein Sericin, which are produced in the larval tissue silk gland (SG) [6]. A transgenic technique has been applied to the silkworm [7], which has enabled the production of a large amount of recombinant proteins through the introduction of transgenes for overexpression in the SG [8]. The silkworm can be utilized as a significant bioreactor by this approach.

Based on its biological and industrial importance, the whole genome sequence of the silkworm was reported in 2004 by two research groups [9, 10]. This was the first lepidopteran genomic analysis and has served as a fundamental basis for genomic studies on Lepidoptera. This silkworm genome data was updated in 2008 [11] and related data have since become available, including microarray-based gene expression profiles, a BAC-based linkage map, and full-length cDNA data [12–14]. These data have strongly promoted studies on *B*. *mori* and other lepidopteran insects in the past few decades.

A new and high-quality reference genome assembly of the silkworm p50T (*daizo*) strain using PacBio long-read and Illumina short-read sequencers was reported in 2019 [15]. The new genome assembly consists of 696 scaffolds with N50 of 16.8 Mb and only 30 gaps. Predicted new gene model based on this novel genome assembly, using cDNA, protein and RNA-seq data as hints, was constructed and was more precise that the previous model made via the old genome assembly [15]. The next step in the establishment of genome-related data is transcriptome data, which contains reference transcriptome sequence data as well as gene expression profiles in major tissues. These data will significantly contribute to advances in research on the silkworm and other lepidopterans.

In the present study, we constructed a reference transcriptome dataset using RNA-seq data obtained from ten major tissues/subparts of silkworm larvae (Fig. 1). RNA-seq data were mapped on the new genome assembly and reference transcriptome sequence data were successfully constructed. We also performed functional annotation of the reference transcriptome using human and *Drosophila* protein datasets in addition to NCBI-nr data. Established reference transcriptome sequence data provided a nearly complete structure for *sericin*-*1* (*ser1*), a major silk gene with a complex sequence. The expression of the transcriptome was investigated in each tissue, and the expression of a number of transcripts was found to be confined to tissues such as testis (TT). Among them, genes with transcripts that were strongly expressed in the midgut (MG) formed tight genomic clusters, suggesting that they originated via gene duplication. The transcripts of transcriptional factor (TF) genes expressed in specific tissues or SG subparts were also detected, and we speculate that these genes play key roles in the major biological process of these tissues/territories. The present results will accelerate molecular biological studies on the silkworm and other related species, and this is an essential milestone to promote entomological research as well as the practical use of insects.

**Fig. 1.**
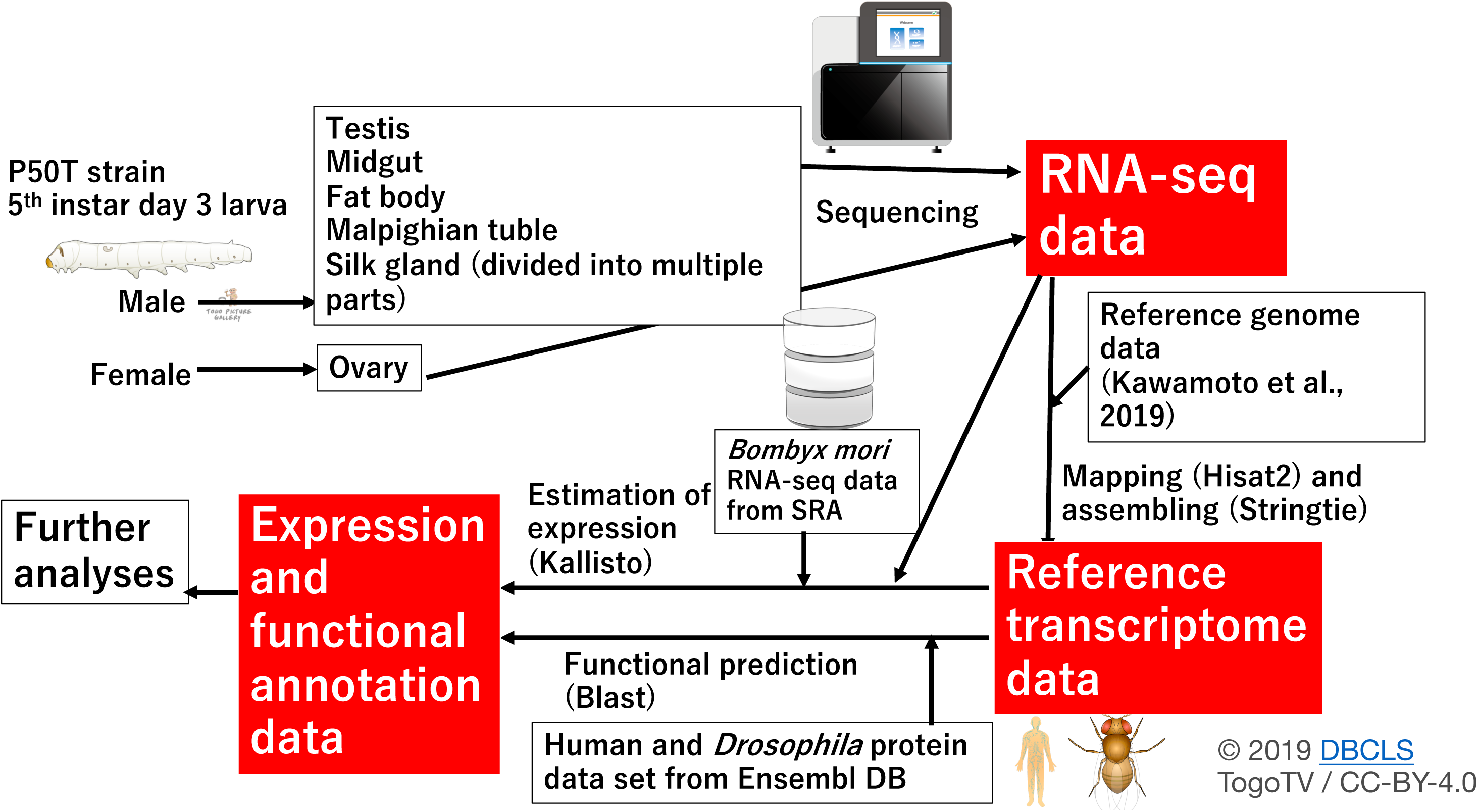
Workflow of the data analysis performed in the present study. To obtain reference transcriptome sequences, Fastq data of 10 tissues/subparts from 5th instar larvae were mapped to the new reference genome [15]. Kallisto software was used to estimate the expression abundance of each transcript in these tissues. Regarding RNA-seq data obtained from the public database (o751 strain), see Table 1. We performed a Blast search against human and *Drosophila* genome data to perform functional annotations of the reference transcriptome. Insect, human, database images, and sequencer drawings [71] are licensed at Creative Commons [72].

**Table 1.**
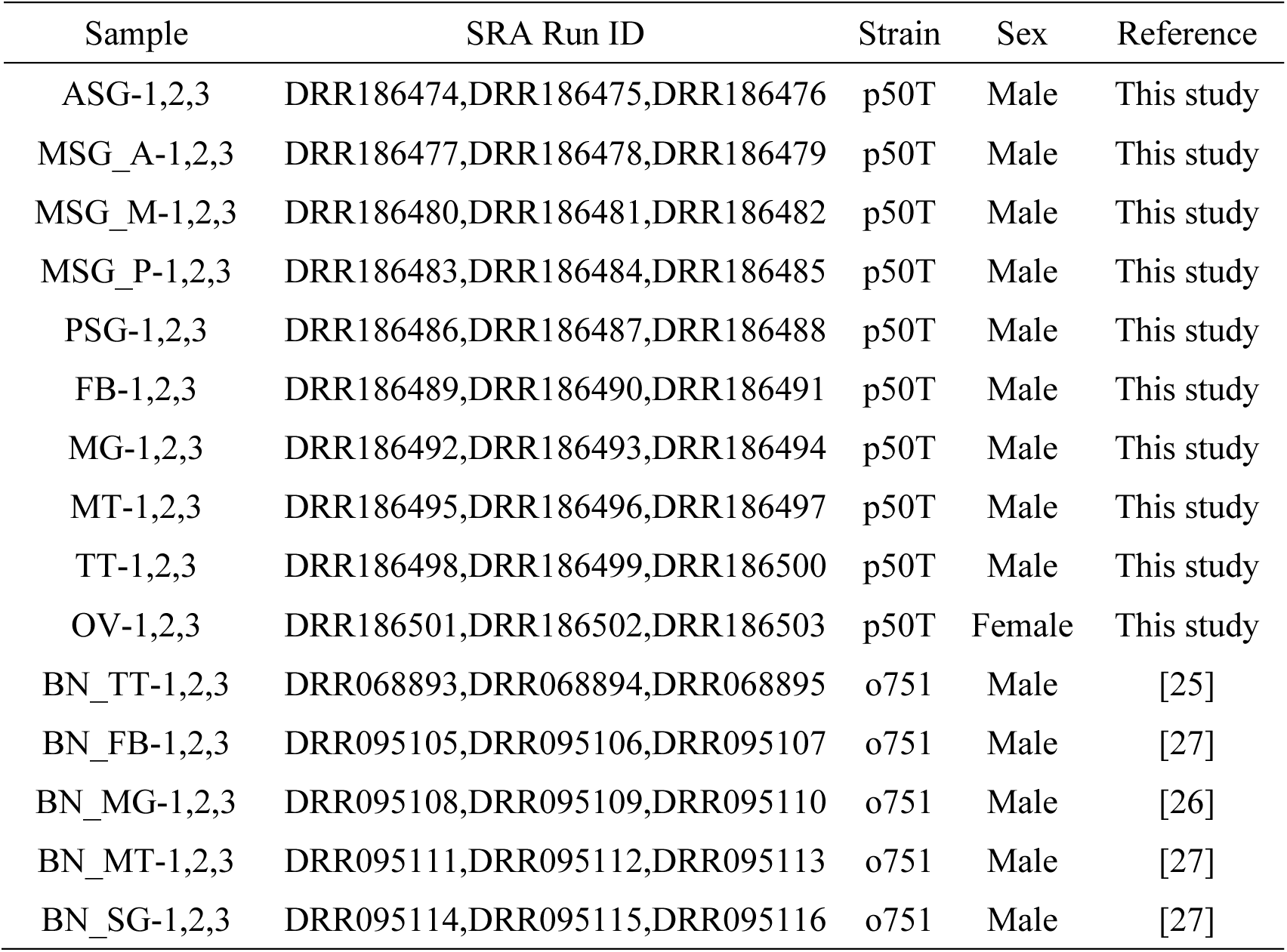
Samples for RNA-seq and run accession IDs.

## Results

### Reference transcriptome data

We performed a transcriptomic analysis of major silkworm larval tissues, namely, the SG, fat body (FB), MG, Malpighian tubules (MT), TT, and ovary (OV), to acquire more expanded RNA-seq data (Fig. 2; Table 1). Gene expression was clearly differentiated among subregions in the SG [16, 17], and, thus, we subdivided the SG into five subparts and investigated gene expression in each region (the anterior silk gland (ASG), anterior part of the middle silk gland (MSG_A), middle part of the middle silk gland (MSG_M), posterior part of the middle silk gland (MSG_P) and posterior silk gland (PSG)) (Fig. 2; Table 1). Totally ten tissues/subparts were dissected from the fifth instar third day larvae of p50T strain with three biological replicates (Table 1). Thirty sets of RNAs were used for RNA-seq. We mapped RNA-seq data on the reference genome assembly using information from the previously established gene model [15] (hereafter referred to as “gene model data (GMD)”) and constructed reference transcriptome data (RTD). RTD comprises 51,926 transcripts in 24,236 loci (Fig. 3A), the numbers of which are higher than those of GMD (24,236 vs 16,845 loci and 51,926 vs 16,880 transcripts; see Fig. 3A and 3B). Therefore, RTD is an extension of GMD. To perform functional annotations, coding sequence (CDS) regions and amino acid sequence data were constructed by using RTD (Additional file 1), and found that 39,619 transcripts, derived from 16,632 loci, had at least one CDS in RTD. The predicted amino acid sequences were used for gene functional annotations by a homology search against human and *Drosophila* gene sets. This analysis revealed that 26,698 genes showed homology to human genes and 29,177 to fruit fly genes (Additional file 2). We also performed a blastp analysis using the NCBI nr database and found that 43,358 amino acids had homological proteins in this database (Additional file 3).

**Fig. 2.**
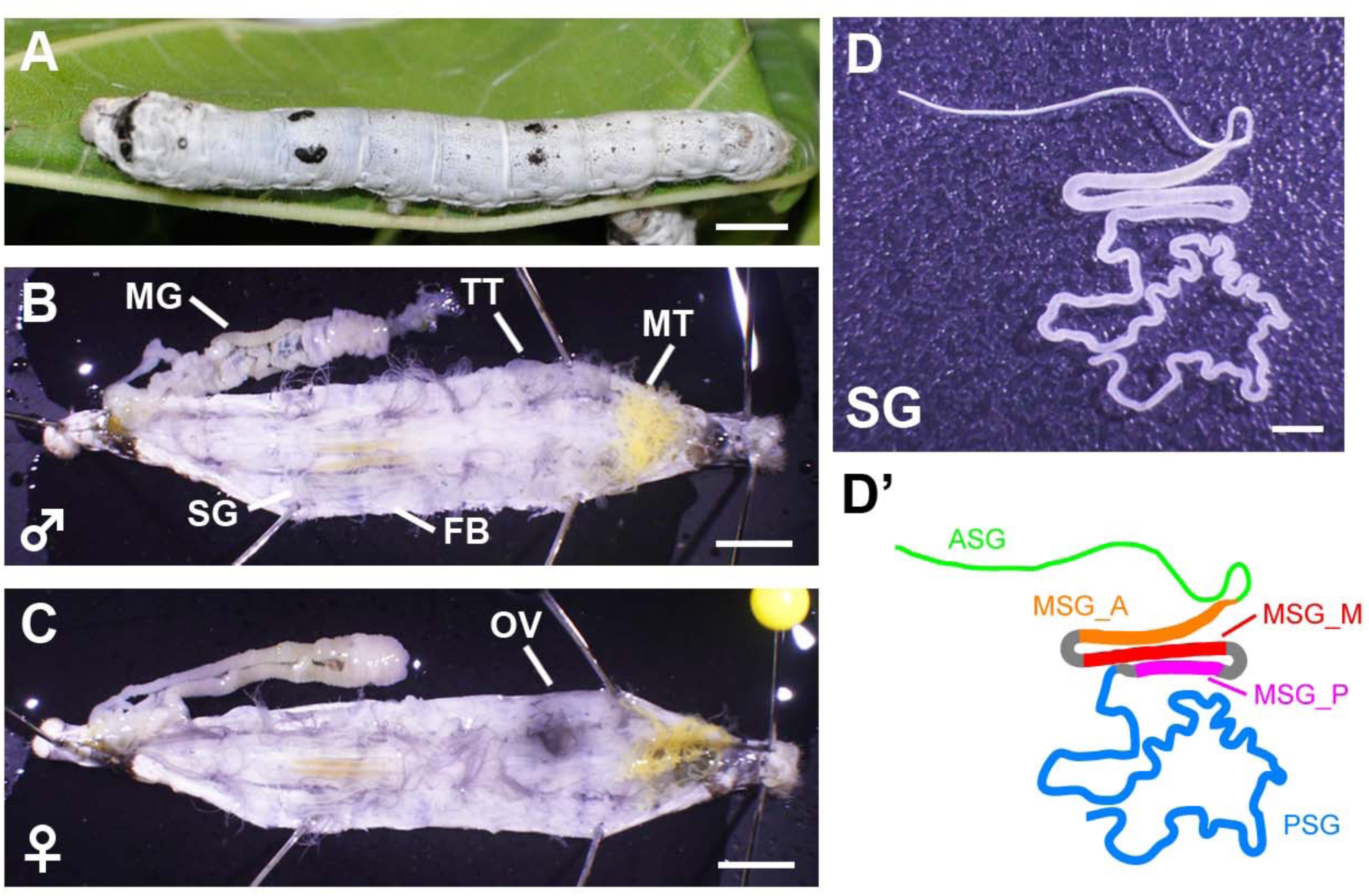
(A) Final (5th) instar larva of the silkworm p50T (*daizo*) strain. Scale bar = 5 mm. (B, C) Male (B) or (C) female individuals dissected on the third day of fifth instar larvae. Scale bar = 5 mm. MG, mid gut; TT, testis; MT, Malpighian tubules; SG, silk gland; FB, fat body; OV, ovary. (D, D’) Image (D) and schematic (D’) of the silk gland. ASG, anterior silk gland; MSG_A, anterior part of the middle silk gland; MSG_M, middle part of the middle silk gland; MSG_P, posterior part of the middle silk gland; PSG, posterior silk gland. Scale bar = 2.5 mm.

**Fig. 3.**
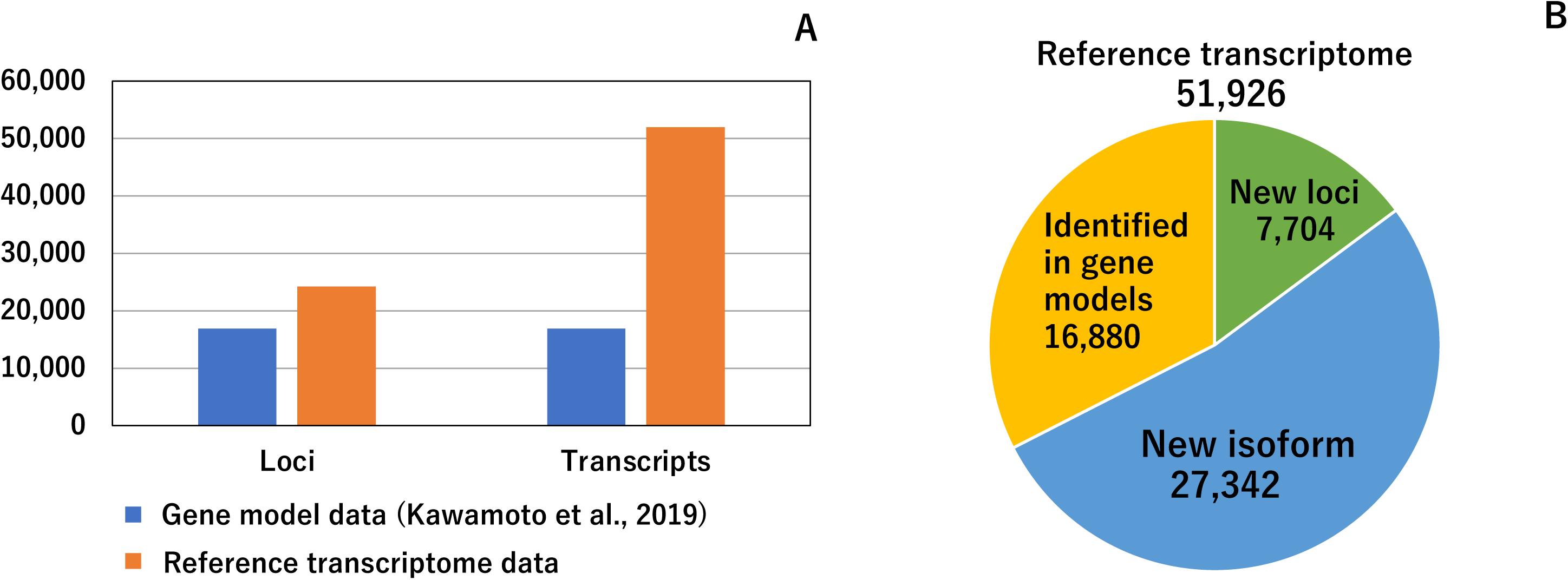
Basal characteristics of the reference transcriptome. (A) Comparison of gene model data [15] and the reference transcriptome data of the present study. The numbers of loci and transcripts are shown. These numbers were calculated from gff files of the two data sets. (B) Classification of 51,926 transcripts. Each transcript was classified into three categories, and the numbers of the three categories are shown in a pie chart. Definitions of the three categories are described in the main text.

### Comparison between constructed reference transcriptome data and previous gene model data

RTD represents a marked improvement over GMD. Several misassembled genes are present in GMD, as represented by KWMTBOMO00087-88, 00196-197, or 00222-223 genes [15]. These genes are split into two structures even though full-length cDNA data define them as single genes [14, 15]. We investigated their structures in our model and found that all were accurately predicted (MSTRG.494.1, MSTRG.649.1-2, and MSTRG.704.1-3) (Additional file 4: Fig. S1). The elucidated gene structure was attributed to the extensive RNA-seq analysis in the present study; the previous study lacked gene expression data from tissues such as TT, OV, MT, and PSG, and these genes were all strongly expressed in these tissues (Additional file 5: Table S1).

RTD also provided a number of novel genes/isoforms. Comparisons of RTD with GMD revealed that among the 51,926 transcripts identified, 7,704 belonged to the new loci group, whereas 27,342 were categorized as new isoforms (Fig. 3B, see Methods). Among the 7,704 new loci group transcripts, a number of genes were also present in the previously established full-length cDNA-based gene model [14]. However, 2,324 transcripts did not hit this gene set and, thus, these genes to which these transcripts belong were perceived to be novel genes. An expression analysis revealed that many of these genes were commonly expressed in all of the tissues investigated herein, whereas other genes were exclusively expressed in specific tissues, such as TT (Additional file 6: Fig. S2). The functional annotation analysis revealed that newly identified genes included a trypsin inhibitor (MSTRG.14562.2), carboxypeptidase (MSTRG.16874.1-3), and pyruvate kinase (MSTRG.18651.2). Therefore, our RTD represents a significant improvement over GMD (Additional file 3).

### Elaborated structures of silk genes

Silk production is one of the most prominent characteristics of the silkworm. Silk genes are strongly expressed in the silk-producing tissue SG and our extensive RNA-seq data are expected to show highly elaborated models for these genes. *ser1* is one of the major silk genes, is strongly expressed in the MSG, and encodes a >400-kDa serine-rich protein [6, 18]. *ser1* is composed of 9 exons, among which exon 6 has a long repetitive sequence with a length of ∼6,500 bp [6, 19]. The full-length sequence of this exon has yet to be elucidated due to its complexity. We herein demonstrated that our model MSTRG.2477.1 provided an almost complete sequence for this exon (6,234 bp, Additional file 7: Fig. S3). Exon 6 encodes serine, glycine, threonine, asparagine, and aspartic acid-rich residues (Additional file 8: Fig. S4), which is consistent with previous findings showing that Ser1 comprises large numbers of these residues (Additional file 9: Table S2) [20]. The detailed structural analysis revealed that the long repetitive motif identified here comprised 53 repeats of a 38-amino acid unit (Additional file 10: Fig. S5). Each unit had serine-rich residues and a slight difference was observed in sequences among units (Additional file 10: Fig. S5) [6]. The 38-amino acid-based repeat unit was also observed in exon 8 of *ser1* (Additional file 8: Fig. S4; Additional file 10: Fig. S5) or in the sericins of saturniid species [6, 21], and, thus, the repeat unit of this length is expected to have a structural function in a number of sericin proteins.

We also observed significant improvements for other *sericin* genes. Sericin-3 (Ser3) is another major silk protein that has a relatively soft texture and possesses serine-rich residues [22, 23]. In GMD, a 73-bp deletion was detected in exon 3, and due to this structural error, a frame shift was present in the predicted amino acid sequence (KWMTBOMO06311; Additional file 11: Fig. S6). In contrast, our RTD (MSTRG.2595.1) successfully provided an accurate gene structure (Additional file 11: Fig. S6). Sericin-4 (Ser4) is another sericin protein that is composed of 34 exons [24]. This gene is split into three distinct structures in GMD (KWMTBOMO06324, KWMTBOMO06325, and KWMTBOMO06326), whereas RTD provided an exact model (MSTRG.2610.1) (Additional file 12: Fig. S7). Collectively, these results suggest that our RTD provided highly defined structures, even for complex silk genes.

### Estimating the abundance of the reference transcriptome in multiple tissues

Our extensive transcriptomic analysis provided fundamental insights into gene expression profiles in multiple silkworm tissues. The expression abundance of each transcript was calculated as transcripts per million (tpm) (Additional file 13) and overall gene expression was compared among tissues through two independent methods, one for hierarchical clustering (HC) and another for a principal component analysis (PCA). To avoid the effects of low expression transcripts, we performed these analyses using transcripts with tpm values > 30 in at least one sample. HC using all transcript tpm data was also performed for comparisons. These analyses revealed that the biological replicate samples collected herein were highly reproducible because they were derived from the same tissues forming tight clusters (Fig. 4; Additional file 14: Fig. S8). In the HC analysis, a single cluster was formed for the MSG_M and MSG_P in each sample (Fig. 4; Additional file 14: Fig. S8B), and we speculate that this was due to highly conserved gene expression between these SG territories. A correlation analysis of gene expression using SG transcriptome data supported this hypothesis (Additional file 15: Table S3). A previous study performed RNA-seq analysis of multiple silkworm larval tissues in another silkworm strain, o751 [25–27], and these RNA-seq data were added to our analysis (Fig. 4; Table 1; Additional file 14: Fig. S8; these samples are referred to as BN_MG, BN_FB, BN_MT, BN_SG, and BN_TT). We found that the samples collected from the same tissues clearly formed clusters (Fig. 4). In the o751 strain, the SG was collected as a whole tissue. Although all the SG subparts (MSG_A, MSG_M, MSG_P, ASG and PSG) were closely located in the HC analysis of transcripts with > 30 (tpm) in at least one sample, BN_SG formed a single cluster distant from those of the SG subparts (Fig. 4). Therefore, our transcriptomic data is a robust platform for analyzing and comparing gene expression in multiple tissues.

**Fig. 4.**
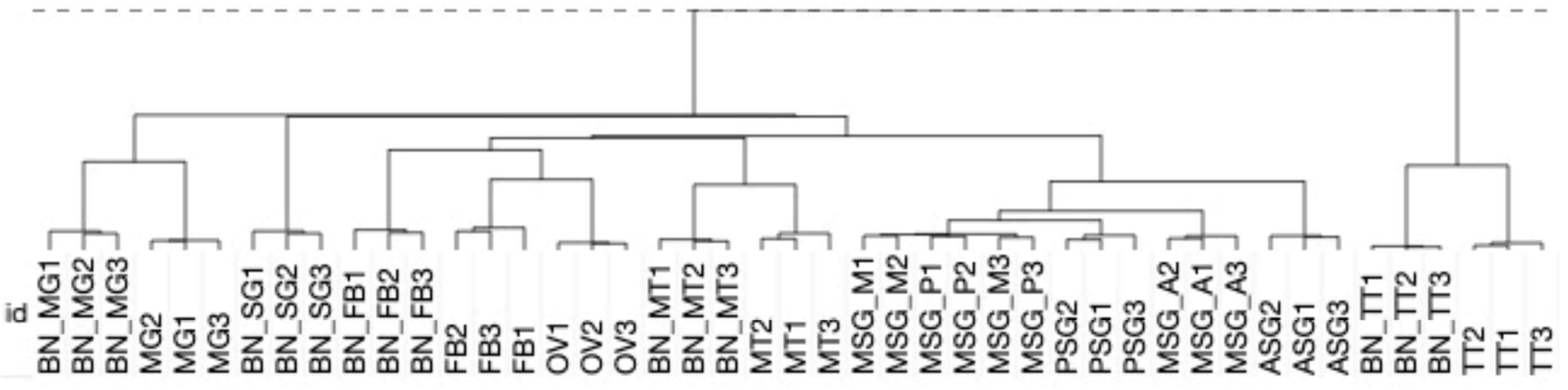
Hierarchical clustering of expression data in 45 samples of transcripts showing a tpm value > 30 in at least one sample. Abbreviations that start with “BN” indicate samples collected in the previous study [25–27]. The numbers added to the abbreviations mean biological replicates.

### Transcript abundance in each tissue and in the silk gland

Using the data described above, we investigated gene expression in detail in each tissue. We focused on genes with a tpm value > 30, which approximately accounted for the top 5% of the most strongly expressed genes (data not shown), and regarded such transcripts as being expressed in each tissue. We found that 711 genes were expressed in all tissues (Fig. 5), suggesting ubiquitous functions. We also detected genes expressed in specific tissues (Fig. 5, Additional file 16: Table S4). Among them, genes solely expressed in TT were the most abundant (1882), followed by those expressed in OV (799), MT (499), and MG (440) (Fig. 5, Additional file 16: Table S4). We also identified genes expressed in more than two tissues, such as TT and OV (397, Fig. 5). A functional enrichment analysis (FEA) was performed on genes with tissue-restricted expression, and functional clusters enriched in each tissue were very diversified; for example, genes exclusively expressed in the TT were strongly enriched for “cilium organization”, “Huntington’s disease”, and “cilium or flagellum-dependent cell motility” (Fig. 6A) whereas “Metabolism of RNA”, “regulation of mRNA metabolic process”, and “ribonucleoprotein complex biogenesis” were enriched in the OV (Fig. 6B). Comparisons of gene expression levels among tissues revealed that the expression levels of strongly expressed genes were very high in the ASG (Fig. 7). These genes comprise fungal protease inhibitors, cuticular protein genes and others (Additional file 17: Table S5). In contrast, the levels of strongly expressed genes were lower in the MSG-M/MSG-P (Fig. 7; Additional file 17: Table S5). We also investigated the expression profiles of genes strongly expressed in each tissue and found tissue-restricted expression for these genes in the MG, FB, and MSG_A and ubiquitous expression for those in the other tissues examined (Additional file 17: Table S5; Additional file 18: Fig. S9). The genomic positions of tissue-enriched genes were examined, and genes strongly expressed in the MG formed tight genomic clusters (Fig. 8; Additional file 17: Table S5). We also found clusters for strongly expressed genes in other tissues (Additional file 17: Table S5; Additional file 19: Fig. S10).

**Fig. 5.**
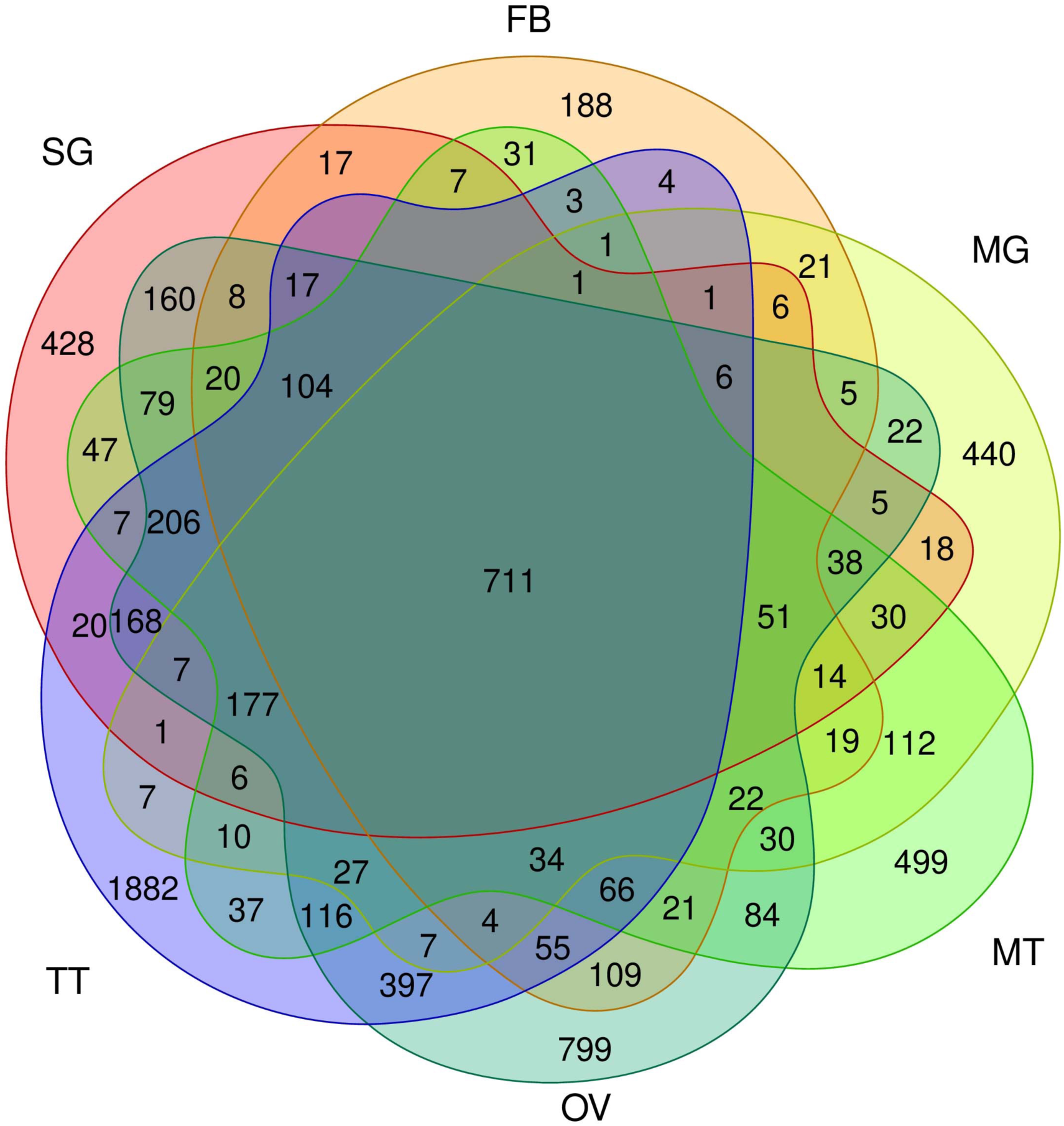
Venn diagram showing transcripts expressed in each tissue. The number of transcripts with a tpm value >30 is shown. SG, silk gland; FB, fat body; MG, mid gut; MT, Malpighian tubules; OV, ovary; TT, testis.

**Fig. 6.**
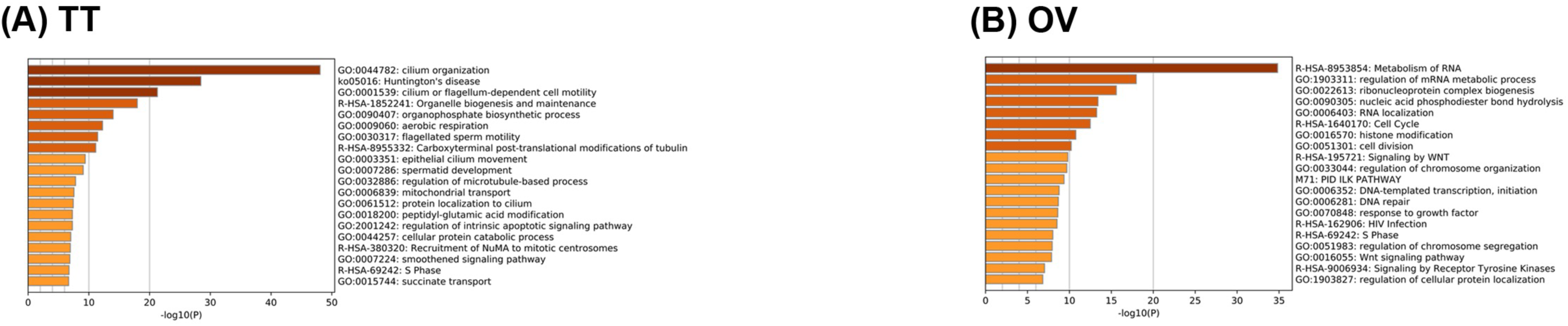
Results of the enrichment analysis by Metascape in testis (TT) (A) and ovary (OV) (B). An enrichment analysis was performed using annotation data against the human gene set of the reference transcripts expressed in specific tissues. −log10 (P) represents −log10 (P-value). For example, −log10 (P)=5 represents P-value= 10^-5^.

**Fig. 7.**
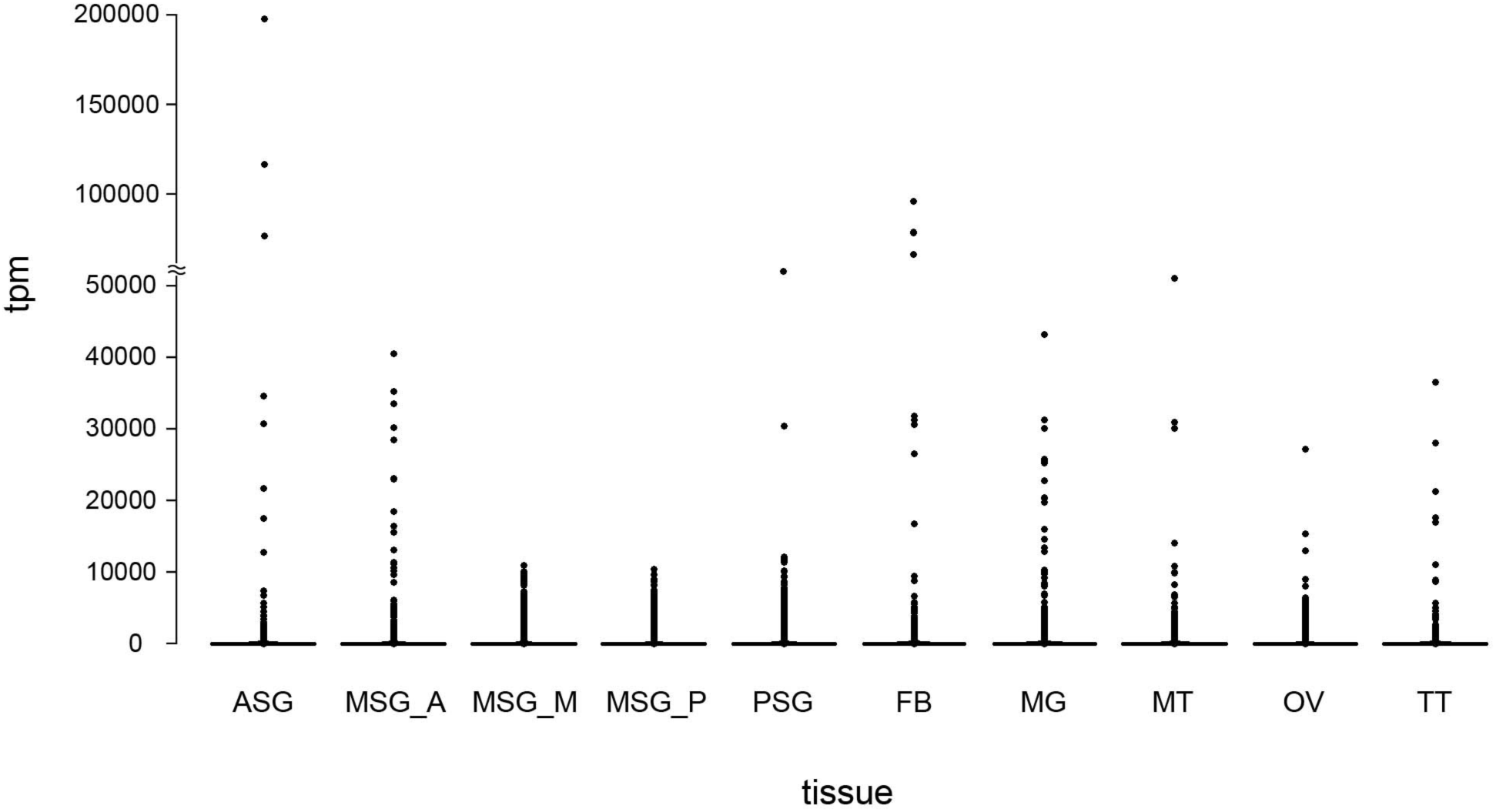
A scatter plot of gene expression in each tissue. Each spot shows the tpm value.

**Fig. 8.**
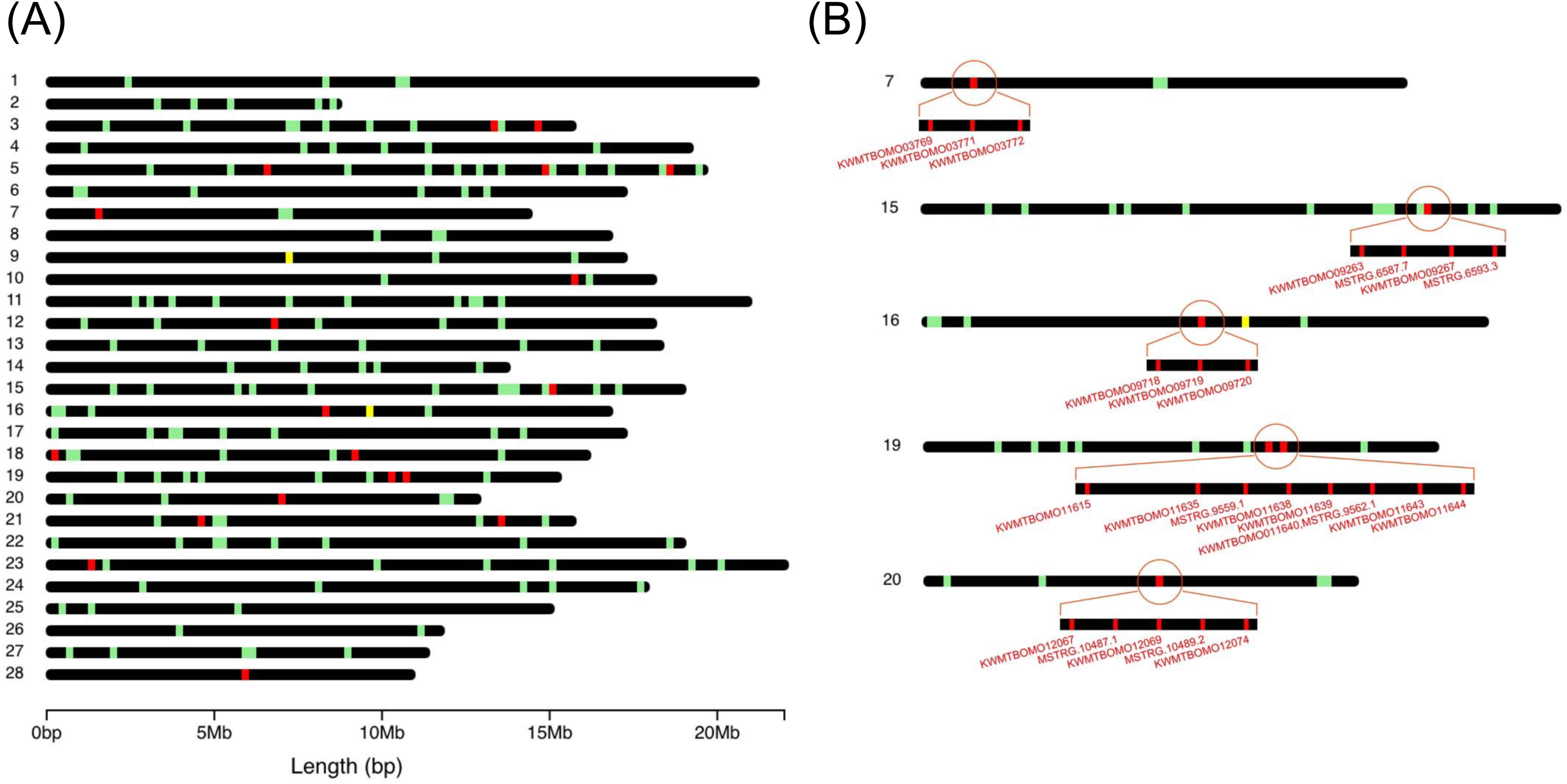
Genomic position of genes strongly expressed in the MG only (red bar), in other than the MG (green bar), and commonly in the MG and other tissues (yellow bar). The top 50 strongly expressed genes are shown. The black bar indicates the chromosome. The number at the left side of each chromosome indicates the chromosomal number. (A) Genomic positions of all chromosomes. (B) Genomic positions of chromosomes 7, 15, 16, 19 and 20, in which the tight clusters of strongly expressed genes in the MG are present.

We then investigated gene expression in the SG in more detail. Previous studies revealed that a number of genes showed territory-specific expression in the SG [16, 17]. However, overall gene expression in each territory remains unclear. We herein demonstrated that >1000 genes were commonly expressed in all SG subparts and also that a number of genes were expressed in specific territories (Fig. 9). They included 351 ASG-restricted, 180 MSG_A-restricted, 99 MSG_M-restricted, 71 MSG_P-restricted, and 100 PSG-restricted genes, respectively (Fig. 9, Additional file 20: Table S6). Furthermore, we identified genes that were commonly expressed in more than two territories (Fig. 9). They included genes expressed in the MSG_M and MSG_P (Fig. 9), and combined with the results of HC and the correlation analysis (Fig. 4; Additional file 15: Table S3), we speculate that gene expression is highly conserved between the MSG_M and MSG_P. This result was supported by the presence of a smaller number of genes expressed solely in the MSG_M or MSG_P (Fig. 9). We also found that the genes that were exclusively expressed in the ASG were more abundant than in other territories, and fewer genes were commonly expressed in the ASG and other subparts (Fig. 9). We speculate that this reflects the functional diversification of the ASG because the numbers of genes exclusively expressed in ASG (351) and those expressed in the MSG and PSG (395) were comparable, suggesting similar diversified functions (Fig. 9). This may also be the case for the MSG_A based on the presence of similar characteristics (Fig. 9). FEA revealed that the functional clusters enriched in each SG subpart were largely diversified (Additional file 21: Fig. S11).

**Fig. 9.**
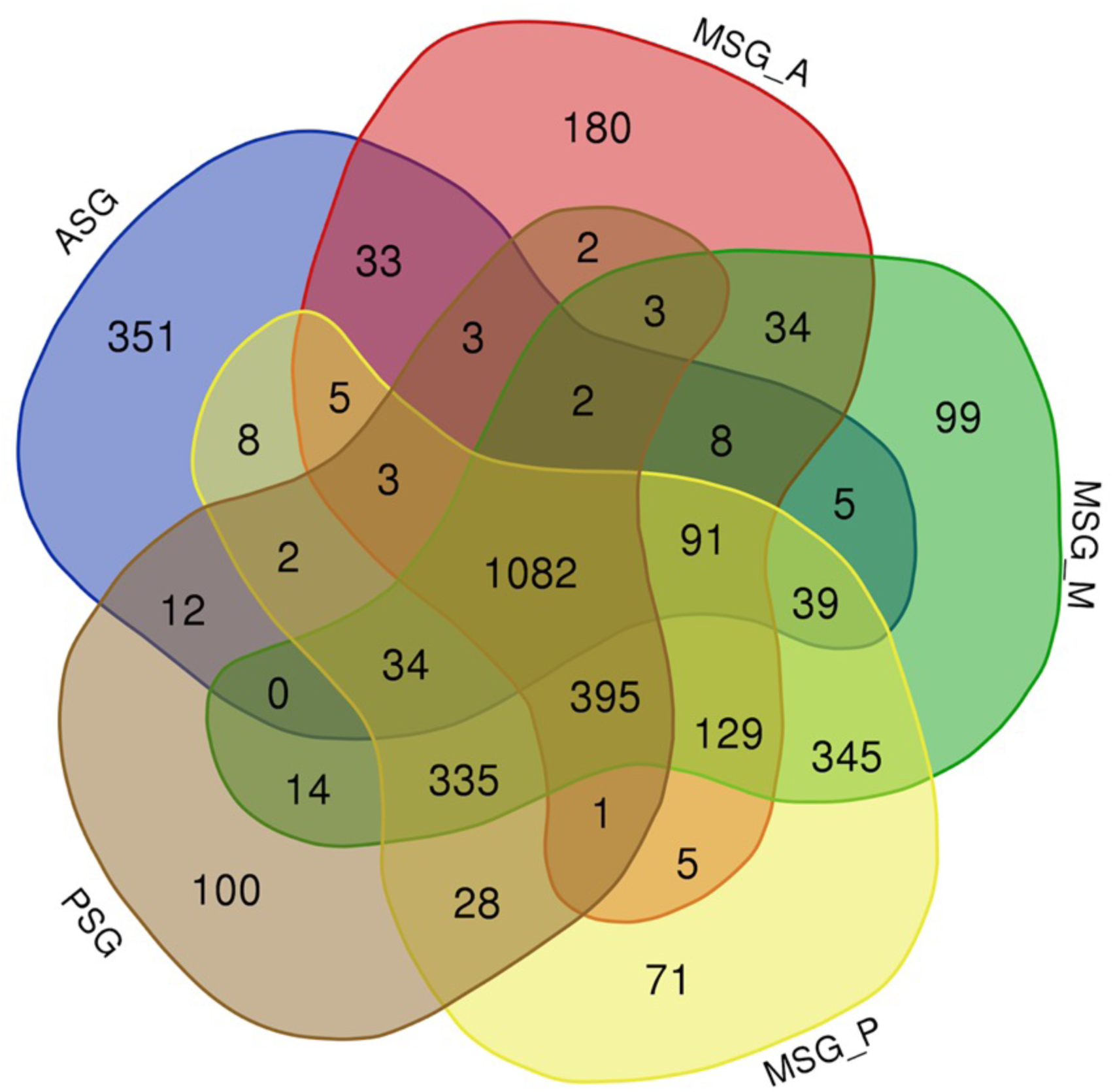
Venn diagram showing the number of transcripts with a tpm value >30 in each silk gland part.

### Expression analysis of transcriptional factor genes in the silkworm

A transcriptomic analysis is a powerful tool for identifying genes with low levels of expression. TF genes are considered to show low expression levels even though they have many important functions in developmental, physiological, and other major biological processes. Therefore, an expression analysis of TF genes will contribute to a more detailed understanding of the silkworm biology. Silkworm TF genes have recently been catalogued [28] and we investigated their expression levels in various silkworm tissues using this information. According to the low level of expression of TF genes, we herein perceived TF genes with a tpm value > 5 as those expressed in each tissue. This analysis revealed that a number of TF genes were exclusively expressed in the OV and/or TT (Fig. 10A; Additional file 22: Table S7). These genes included KWMTBOMO02002 (*traffic jam*), KWMTBOMO002212 (*mirror*), KWMTBOMO01693 (*vismay*), and KWMTBOMO06584 (*Sox100B*), all of which play significant roles in gonad morphogenesis, oogenesis, spermatogenesis, and TT differentiation in *Drosophila melanogaster* (Additional file 22: Table S7) [29–32]. We also found that KWMTBOMO09369/KWMTBOMO10218, the silkworm counterparts of human GATA4, were expressed in the OV, similar to that in humans (Additional file 22: Table S7) [33]. We speculate that these TF genes have conserved functions in a wide variety of organisms, possibly in the development, differentiation, or homeostasis of reproductive tissues.

**Fig. 10.**
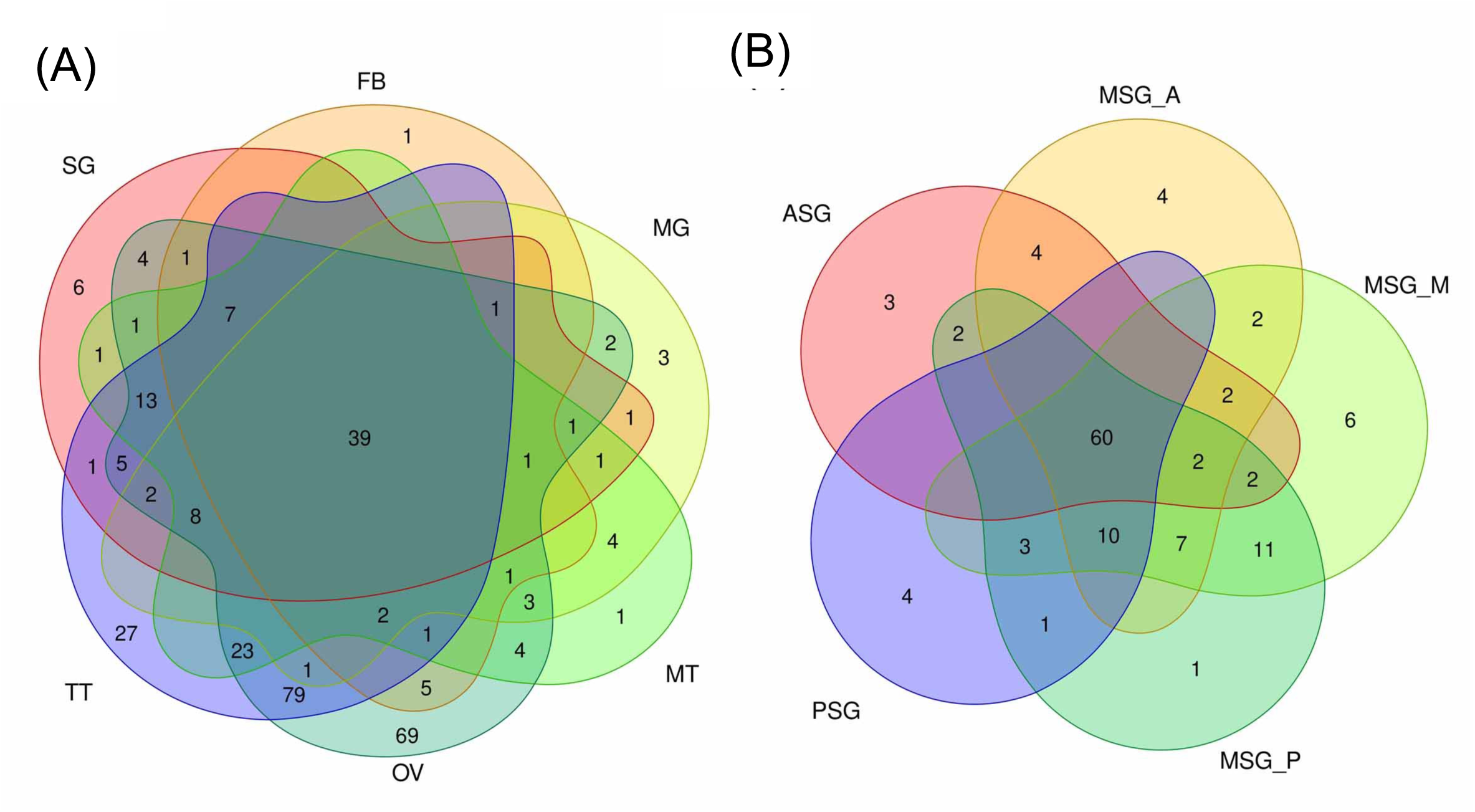
Venn diagrams showing the number of TF genes with a tpm value >5 in each tissue (A) and in silk gland subparts (B).

We also investigated TF gene expression in the SG. Previous studies revealed the central roles of the TF genes *forkhead* (*fkh*), *Antennapedia* (*Antp*), and *Arrowhead* (*Awh*) in the regulation of silk gene expression [34–38]. Other TF genes that function in the regulation of silk gene expression may also exist because silk genes are strongly expressed and a number of silk genes are expressed in specific territories in the SG (Fig. 9). In addition to previously identified TF genes, the expression of novel TF genes, such as those belonging to the High Mobility Group (HMG), Zinc Finger (ZF), Paired box (PAX), or forkhead family, was confined to the SG (Fig. 10B, Additional file 23: Table S8). The tpm values of these genes are shown in Additional file 23: Table S8, and we identified TF genes expressed solely in the ASG or MSG_A. Genes expressed in more than two subparts were also detected (Additional file 23: Table S8). We confirmed their expression in an RT-PCR analysis and experimental results were consistent with tpm values (Additional file 24: Fig. S12). The TF genes identified herein include those with indispensable roles in the development of *D*. *melanogaster*; for example, *Dichaete* (*D*), *gooseberry-neuro* (*gsb-n*), and *sloppy paired 2* (*slp2*) are essential for embryonic segmentation [39–41], *Enhancer of split mβ-HLH* (*E*(*spl*)*mβ-HLH*) for neurogenesis [42], and *twin of eyegone* (*toe*) for eye development [43]. We consider that it is very interesting if these genes have significant roles in the regulation of the silk gene expression, a function that is independent from the developmental regulation, in the silkworm.

## Discussion

In the present study, we performed RNA-seq analysis of multiple larval tissues from the silkworm *B*. *mori*. We established RTD using a recently reported high-quality reference genome assembly [15] and RNA-seq data newly obtained herein. RTD showed marked improvements over GMD, most notably the establishment of a nearly complete structure for *ser1* (Additional file 7: Fig. S3; Additional file 8: Fig. S4); its full sequence has never been entirely elucidated due to its complexity. Our results indicate that an extensive RNA-seq analysis in combination with high-quality reference genome data provides highly refined gene structures, even for complex genes. The cost of performing a deep sequencing analysis has recently decreased and, thus, it has become affordable for every researcher to conduct not only a short-read RNA-seq analysis, but also long-read genome sequencing using their own species. The present results are a significant proof-of-concept that highly refined gene structures may be established using the combination of these data, even for non-model organisms. Furthermore, more elaborate gene structures may be constructed using the RNA-seq data derived from other tissues and/or stages.

We herein found that a number of genes showed tissue-restricted expression in the silkworm (Fig. 5). Among them, genes exclusively expressed in the TT were the most abundant (Fig. 5). A previous study identified a number of TT-specific genes in the silkworm [12], which is consistent with the present results. The presence of a number of TT-specific genes was also demonstrated in the jewel wasp, *Nasonia vitripennis* [44], indicating that this is a common feature in insects. Recent studies on *Drosophila* revealed that newly emerging genes were strongly biased for expression in the male reproductive system [45]. Therefore, the TT-specific genes identified in the present study may have similar traits. This issue may be confirmed by investigating the evolutionary ages of these genes, and, if this is the case, addressing the question of why the TT is a tissue that is permissive for new gene birth, a phenomenon observed not only in insects, but also in vertebrates [46], which will become possible using the silkworm. Another important result obtained from our cross-tissue gene expression analysis is that genes strongly expressed in the MG showed strong tissue-restricted expression and also formed tight genomic clusters (Fig. 8; Additional file 17: Table S5; Additional file 18: Fig. S9). Comparisons of the sequences of these genes revealed that genes in each cluster encoded homological proteins; chr.7 genes encoded trypsins, chr.15 juvenile hormone-binding proteins (JHBPs), chr.16 fatty acid-binding proteins, chr.19 actin cytoskeleton-regulatory complex proteins, and chr.20 multiprotein bridging factor 2 (MBF2) (Additional file 17: Table S5). Among these genes, strong expression in the MG has already been demonstrated for *jhbp*s [47], and Trypsin proteins were present in digestive juices [48]. Homological genes that cluster in the genome are generally considered to have originated via tandem gene duplication, and to the best of our knowledge, few studies have investigated clustered genes that are expressed in the MG. In one case study on *Drosophila*, neutral lipase genes expressed in the MG clustered in the genome and were presumably under positive selection to retain different substrate specificities towards new lipid components of the diet [49].

Based on these findings, the genes identified in the present study may also have advantages in the silkworm MG, such as enhancing activity for digestion and/or xenobiotic detoxification. The present study provides valuable insights into gene evolution as well as neofunctionalization in insects, which may be validated in more detail in future studies. In addition, the results obtained herein will facilitate the practical application of the silkworm; targeted gene integration into the clusters identified in the present study will enable strong gene expression in the MG, which will contribute to the establishment of strains with valuable properties, such as increased antiviral or antibiotic activities. The latest genome editing technologies should promote the establishment of useful silkworm strains.

Our detailed gene expression analysis provides fundamental information on the traits of the SG, particularly each subpart. The SG is a tissue that arises from a single embryonic segment and has a long tubular structure [50]. Gene expression in each SG subpart is largely diversified, as demonstrated herein (Fig. 9; Additional file 20: Table S6) and in previous studies [16, 17]. The most important result of the present study is that overall gene expression in the ASG was diversified the most within the SG, as demonstrated by the presence of a number of genes exclusively expressed in the ASG (351, Fig. 9), a number of genes expressed both in the MSG and PSG (395, Fig. 9), and the location of the ASG at the outermost site in the SG cluster in the HC analysis (Fig. 4). We speculate that these results indicate the functional diversification of the ASG, which is consistent with previous findings showing that the ASG functions in silk fiber processing and the MSG/PSG in silk protein production [51]. FEA revealed that functional clusters enriched in the ASG were “carbohydrate metabolic process” and “Transport of small molecules” (Additional file 21: Fig. S11A), while those enriched in the MSG/PSG were “ribonucleoprotein complex biogenesis” and “Translation” (Additional file 21: Fig. S11F), further supporting this concept. In this context, another result showing that gene expression in the MSG_A also showed diversification is of interest. We demonstrated that genes specifically expressed in the MSG_A were abundant (180, Fig. 9), a number of genes were commonly expressed in the MSG_M/MSG_P/PSG (335, Fig. 9), and the MSG_A was located at the outermost site in the MSG/PSG cluster in the HC analysis (Fig. 4). MSG_A is a part of the MSG and functions in the production of the sericin proteins Ser2 and Ser3 [23, 52], similar to the function of the MSG_M/MSG_P in the production of Ser1 [20]. Nevertheless, our results indicate that gene expression in the MSG_A was more diverse than that in the MSG_M/MSG_P compared with that of the PSG; the PSG is the territory that produces fibroin and not sericin [53], whereas gene expression appeared to be more conserved between the PSG and MSG_M/MSG_P than between the MSG_A and MSG_M/MSG_P (Figs. 4 and 9). These results may be attributed to the presence of a number of genes that are strongly and specifically expressed in the MSG_A, including ecdysone oxidase, fatty acid hydroperoxide dehydratase, and other genes (Fig. 9; Additional file 20: Table S6). We found that genes strongly expressed in the MSG_A were enriched for the functional clusters of “Metabolism of vitamins and cofactors” and “organic hydroxy compound transport” (Additional file 21: Fig. S11B), and speculated that these clusters define the biological functions that are unique to this territory. Therefore, our extensive RNA-seq analysis provides fundamental insights into the functions of the SG as well as its evolution, which has not yet been elucidated in detail. In the Lepidoptera, the morphology of the SG is largely diversified among species [54] and the Saturniidae, a family that is phylogenetically close to the Bombycidae, have MSGs with one territory and no morphological separation [6]. Therefore, the differentiation of MSG gene expression observed herein may be specific to *B*. *mori* and/or other closely related species. Further studies are needed to clarify whether differences in gene expression among species are a driving force that generates diversity in cocoon properties, including shape, size, and physical activity. The TF genes identified in the present study may be one of the key factors responsible for the differences in gene expression within the SG or among species.

In the present study, we established RTD for the silkworm and performed a detailed examination of gene expression in silkworm tissues. The results obtained will contribute to our understanding of silkworm biology and further promote the industrial application of the silkworm and other insects.

## Conclusions

We performed RNA-seq analysis of the major larval tissues of the silkworm and established RTD. Using these data, we successfully improved the gene structure and clarified gene expression in detail in each tissue. The present results are a fundamental basis for the further promotion of silkworm research and will contribute to the practical application of insects.

## Methods

### Silkworm rearing, RNA extraction, and sequencing

The silkworm p50T strain was reared on an artificial diet (Nihon Nosan Kogyo, Yokohama, Japan) at 25°C under a 12-hour light/dark photoperiod. The SG, FB, MG, MT, TT, and OV were dissected on the third day of the fifth instar larvae. The SG was further subdivided into the ASG, MSG_A, MSG_M, MSG_P, and PSG. Each tissue/subpart was dissected from one individual, and three biological replicates were obtained and separately analyzed (see Table 1). Tissues were homogenized using ISOGEN (NIPPON GENE, Tokyo, Japan) and the SV Total RNA Isolation System (Promega, Madison, WI) was used for RNA extraction. The total RNA samples extracted were sequenced by Illumina NovaSeq6000 (Macrogen Japan Corp., Kyoto, Japan).

### Construction of RTD and estimation of the expression of each transcript

The raw RNA-seq data of 30 samples were trimmed by Trimmomatic version 0.36 [55]. The trimmed RNA-seq data of each tissue were mapped to the new reference genome with a new gene model [15] by HISAT2 version 2.1.0 [56]. Mapped data were each assembled to transcriptome data by StringTie version 1.3.3 [57]. The 30 transcriptome datasets were merged into one transcriptome dataset, referred to as “a reference transcriptome” by StringTie. GffCompare version 0.10.6 was used [58] for comparisons with the reference transcriptome and previously reported gene sets [15]. The transcripts detected at the newly identified loci were categorized into the “New loci” group, those newly detected at the previously identified loci into the “New isoform” group, and other genes and transcripts into “Identified in gene models” (see Fig. 3B).

To estimate the expression of the reference transcriptome in 30 samples, the raw fastq data of each sample and reference transcript data were used with Kallisto version 0.44.0 [59]. The raw RNA-seq data of multiple tissues in *B. mori* strain o751 from the Sequence Read Archive (SRA) and reference transcript data were used in comparisons of transcriptome data: the accession numbers of raw RNA-seq data are DRA005094, DRA005878, and DRA005094 [25–27].

We used TIBCO Spotfire Desktop (version 7.6.0) software with the “Better World” program license (TIBCO, Inc., Palo Alto, CA; [60]) for the classification of differentially expressed samples in silkworm tissues in HC using Ward’s method. Morpheus was also used [61] for HC. R (version 3.6.0) was used in the PCA analysis [62]. Regarding Venn diagram construction and the scatter plot analysis, R (version 4.0.2) was used [63]. The relationships among gene expression profiles in the SG territories were evaluated using Spearman’s rank correlation.

### Annotation for the reference transcriptome and FEA

Transcoder (version 5.5.0) was used to identify coding regions within transcript sequences and convert transcript sequences to amino acid sequences [64]. Transcriptome sequence sets were compared at the predicted amino acid sequence level by the successive execution of the blastp program in the NCBI BLAST software package (version 2.9.0+) with default parameters and an e-value cut-off of 1e-10 [65]. Regarding the reference database sets to be blasted, human and fruit fly (*D. melanogaster*) protein datasets in the Ensembl database (v.97) were used because the sequences of these organisms were functionally well-annotated and amenable to computational methods, such as a pathway analysis [66]. The names of the top-hit genes in the human and fruit fly datasets were annotated to *B. mori* transcripts utilizing Ensembl Biomart [67] and Spotfire Desktop software under TIBCO Spotfire’s “Better World” program license (TIBCO Software, Inc., Palo Alto, CA, USA) [60]. Functional enrichment analyses were performed using the metascape portal site [68] with annotation results against the human gene set.

### Comparison of gene structures among different models

Gene structures among RTD, GMD, and cDNA-based data were compared in the silkbase [69] or KAIKObase [70]. Amino acid sequences were aligned using CLC Genomics Workbench 20.0.04 (QIAGEN, Aarhus, Denmark).

### RT-PCR

cDNA was synthesized using Superscript IV (Thermo Fisher Scientific Inc., Waltham, MA, USA) according to the manufacturer’s instructions. Five hundred nanograms of total RNAs extracted from the ASG, MSG_A, MSG_M, MSG_P, and PSG were used for the cDNA synthesis. KOD FX neo polymerase (Toyobo, Osaka, Japan) was used for RT-PCR. PCR conditions were as follows: 95°C for 1 min followed by 22 cycles (for *rp49*) or 30 cycles (for TF genes) of 95°C for 30 sec, 58°C for 30 sec, followed by 68°C for 1 min, and additional 68 °C for 1 min after the cyclic phase. Primer sequences are listed in Additional file 25: Table S9.

## Declarations

### Availability of data and materials

The RNA-seq reads supporting the conclusions of the present study are available in the SRA with accession number DRA008737 (The accession number of RNA-seq data of each sample is shown in Table 1).

Reference transcriptome data is available at the Transcriptome Shotgun Assembly (TSA) database under accession IDs ICPK01000001-ICPK01051926, and the Gff file of the transcriptome was available via DOI: 10.18908/lsdba.nbdc02443-001.V001.

The estimated abundance of transcripts is available from the Gene Expression Archive (GEA) in DDBJ under accession ID E-GEAD-315 or from The Life Science Database Archive (DOI: 10.18908/lsdba.nbdc02443-002.V001).

Additional data are available in The Life Science Database Archive (the title in the Archive is “KAIKO - Metadata of reference transcriptome data” (DOI:10.18908/lsdba.nbdc02443-000.V001)) and figshare (DOI: 10.6084/m9.figshare.c.5333894).

#### Additional file 1

Predicted amino acid sequences of reference transcriptome. DOI:10.18908/lsdba.nbdc02443-004

#### Additional file 2

Functional annotations of the reference transcriptome (blast against human and *Drosophila* gene sets). DOI:10.18908/lsdba.nbdc02443-003

#### Additional file 3

Functional annotations of the reference transcriptome (blast against the NCBI nr database). DOI:10.6084/m9.figshare.14192741.

#### Additional file 4: Fig. S1

Comparison of gene structures among gene model data, cDNA-based data, and reference transcriptomic data. (A) Locus around KWMTBOMO00087, (B) KWMTBOMO00196, and (C) KWMTBOMO00222. GMD; gene model data, CBD; cDNA-based data, RTD; reference transcriptomic data. DOI:10.6084/m9.figshare.14205785

#### Additional file 5: Table S1

Tpm values in MSTRG.494.1, MSTRG.649.1-2 and MSTRG.704.1-3. DOI: 10.6084/m9.figshare.14206034

#### Additional file 6: Fig. S2

Expression of new loci transcripts that did not hit the cDNA-based gene model. Numbers indicate transcripts with a tpm value >0.01 in each tissue. SG, silk gland (using average tpm values in the five SG subparts); FB, fat body; MG, mid gut; MT, Malpighian tubules; OV, ovary; TT, testis. DOI: 10.6084/m9.figshare.14217335

#### Additional file 7: Fig. S3

Structure of the *sericin-1* gene. MSTRG.2477.1 has a long exon 6.GMD; gene model data, CBD; cDNA-based data, RTD; reference transcriptomic data. DOI: 10.6084/m9.figshare.14206166

#### Additional file 8: Fig. S4

The presumptive full-length amino acid sequence of sericin-1 deduced by reference transcriptomic data (MSTRG.2477.1). The orange characters show the amino acids encoded by exon 6 and blue characters by exon 8. DOI: 10.6084/m9.figshare.14206736

#### Additional file 9: Table S2

Comparisons of the sericin-1 amino acid composition elucidated by an amino acid analysis and gene model. Mole% is shown. DOI: 10.6084/m9.figshare.14206748

#### Additional file 10: Fig. S5

Sequence of the 38-amino acid-based repeat unit encoded by exon 6 and exon 8 in sericin-1. Exon 6 comprises 53 repeats and exon 8 13 repeats. DOI: 10.6084/m9.figshare.14212673

#### Additional file 11: Fig. S6

Comparison of the sericin-3 sequence among different gene models. The amino acid sequences identified in the previous study (NM_001114644; [23]), derived from cDNA-based data (BMgn014348), reference transcriptomic data (MSTRG.2595.1), and gene model data (KWMTBOMO06311), are shown. Amino acids that differ in KWMTBOMO06311 are shown as white letters. Note that a frame shift occurs in the gene model data. DOI: 10.6084/m9.figshare.14216927

#### Additional file 12: Fig. S7

Comparison of the *sericin-4* gene structure. The gene structures elucidated by the previous study (*sericin-4*; [24]) and modeled by gene model data (GMD) and reference transcriptomic data (RTD) are shown. DOI: 10.6084/m9.figshare.14216951

#### Additional file 13

Expression data of each transcript in multiple tissues. DOI: 10.18908/lsdba.nbdc02443-002.V001

#### Additional file 14: Fig. S8

(A) Principal Component Analysis (PCA) results with expression profiles in 45 samples of transcripts showing a tpm value > 30 in at least one sample. Abbreviations and the numbers of samples are the same as in Fig. 4. The X axis and Y axis are principal components 1 (PC1) and PC2, respectively. (B) Hierarchical clustering of expression data in 45 samples using all transcript tpm values. DOI: 10.6084/m9.figshare.14216993.

#### Additional file 15: Table S3

Spearman’s rank correlation coefficient among silk gland territories. DOI: 10.6084/m9.figshare.14217071

#### Additional file 16: Table S4

List of genes expressed in specific tissues. DOI: 10.6084/m9.figshare.14217206

#### Additional file 17: Table S5

List of genes strongly expressed in each tissue or in silk gland subparts. The top 50 strongly expressed genes are shown. DOI: 10.6084/m9.figshare.14217218

#### Additional file 18: Fig. S9

A scatter plot of gene expression in each tissue. Each spot shows the tpm value. The top ten ranked strongly expressed genes in (A) ASG, (B) MSG_A, (C) MSG_M, (D) MSG_P, (E) PSG, (F) FB, (G) MG, (H) MT, (I) OV and (J) TT are marked in red. DOI: 10.6084/m9.figshare.14217227

#### Additional file 19: Fig. S10

Genomic position of genes strongly expressed in each tissue. In (A), genes strongly expressed in the ASG only are shown with a red bar, in subparts other than the ASG with a green bar, and those commonly expressed in the ASG and other tissues with a yellow bar. This same applies to MSG_A (B), MSG_M (C), MSG_P (D), PSG (E), FB (F), MT (G), OV (T), and TT (I). (A’) shows the chr. 11 of the ASG and (F’) shows chr. 20 of the FB. Regarding the MG, see Fig. 8. The top 50 strongly expressed genes are shown. The black bar indicates the chromosome, and the number at the left side of each chromosome indicates the chromosomal number. DOI: 10.6084/m9.figshare.14217233

#### Additional file 20: Table S6

List of territory-specific genes in the silk gland. DOI: 0.6084/m9.figshare.14217242

#### Additional file 21: Fig. S11

Results of the enrichment analysis for territory-specific transcripts in the silk gland. The number of transcripts used for the analysis is shown in the bracket. −log10 (P) represents −log10 (P-value). For example, −log10 (P)=5 represents P-value= 10^-5^. DOI: 10.6084/m9.figshare.14217257

#### Additional file 22: Table S7

List of tissue-specific TF genes. DOI: 10.6084/m9.figshare.14217272

#### Additional file 23: Table S8

List of territory-specific TF genes in the silk gland. DOI: 10.6084/m9.figshare.14217296

#### Additional file 24: Fig. S12

RT-PCR of TF genes showing territory-specific expression in the SG. DOI: 10.6084/m9.figshare.14217299

#### Additional file 25: Table S9

Primer sequences used for RT-PCR. DOI: 10.6084/m9.figshare.14217308

## Competing interests

The authors declare no conflicts of interest.

## Funding

This work was supported by the National Bioscience Database Center of the Japan Science and Technology Agency (JST) to HB.

This work was supported by the Cabinet Office, Government of Japan, Cross-ministerial Strategic Innovation Promotion Program (SIP), “Technologies for Smart Bio-industry and Agriculture” (funding agency: Bio-oriented Technology Research Advancement Institution, NARO) to KY, TT, and HS.

This work was also supported by a grant from the Ministry of Agriculture, Forestry and Fisheries of Japan (Research Project for Sericultural Revolution) to KY, TT, and HS.

## Author contributions

Conceived and designed experiments: KY, TT, and HS.

Performed experiments: TT.

Contributed reagents/materials/analysis tools: HS.

Analyzed data: KY, TT, AJ, and HB.

Contributed to the writing of the manuscript under the draft version: KY, TT, and HB.

All authors discussed the data and helped with manuscript preparation. KY supervised the project.

All authors read and approved the final manuscript.

## Acknowledgments

The computing resource was partly provided by the super computer system at the National Institute of Genetics (NIG), Research Organization of Information and Systems (ROIS), Japan. The authors thank Mrs. Kaoru Nakamura and Toshihiko Misawa for rearing the silkworms, Ms. Satoko Kawamoto for technical assistance, and Dr. Jianqiang Sun for data analysis advice.

## Notes

### Competing Interest Statement

The authors have declared no competing interest.

### Summary of Updates

Several analysis were added to the new version.

